# SurvBal: Compositional Microbiome Balances for Survival Outcomes

**DOI:** 10.1101/2023.09.22.559069

**Authors:** Ying Li, Teresa Lee, Kai Marin, Xing Hua, Sujatha Srinivasan, David N. Fredricks, John R. Lee, Wodan Ling

## Abstract

Identification of balances of bacterial taxa in relation to continuous and dichotomous outcomes is an increasingly frequent analytic objective in microbiome profiling experiments. SurvBal enables the selection of balances in relation to censored survival or time-to-event outcomes which are of considerable interest in many biomedical studies. The most commonly used survival models – the Cox proportional hazards and parametric survival models are included in the package, which are used in combination with step-wise selection procedures to identify the optimal associated balance of microbiome, i.e., the ratio of the geometric means of two groups of taxa’s relative abundances.

**Availability and implementation:** SurvBal is available as an R package and a Shiny app: https://github.com/yinglia/SurvBal, https://yinglistats.shinyapps.io/shinyapp-survbal/.

## 1 Introduction

Compositional balances have served as a powerful strategy for implicating bacterial taxa in relation to a wide range of outcomes. The underlying principle of the global balances is that outcomes depend on the ratio of two sets of bacterial relative abundances. Thus, Selbal [1] chooses to identify the (log) ratio of the geometric means of two sets of taxa. Essentially, the numerator of the ratio is the geometric mean of the taxa positively correlated with the outcome while the denominator is the geometric mean of the taxa negatively correlated with the outcome. Philosophically, the global balances include better characterization of the idea of dysbiosis as well as better accommodation of the issue of compositionality which creates significant challenges for interpretation. Operationally, the optimal balances can be identified through greedy search procedures. The approach has been successfully used to identify balances, and the comprising taxa, related to many different outcomes including COVID severity [2], neutrophil levels in HIV infection [3], among others.

However, a limitation in the field is the lack of software for identifying balances in relation to censored survival or time-to-event outcomes. Yet, such outcomes are of considerable interest, particularly as human microbiome studies interface with clinical studies in which time-to-event outcomes (e.g. overall survival, time to relapse, time to disease onset, etc.) are the most commonly used outcomes. Such studies are often subject to significant censoring such that specialized survival models are needed.

We present SurvBal, a flexible R package that facilitates the analysis of balances with censored time-to-event outcomes. It identifies the log-ratio of geometric means of two sets of taxa that is most associated with the survival outcome using a greed step-wise selection approach. The software supports the Cox proportional hazards model and parametric survival models (and by extension, accelerated failure time models), and reports a selected global balance of bacteria increasing vs. decreasing the hazard or survival time. A comprehensive Shiny app is provided for broader users to interactively explore the analytical tool.

## 2 Software Description

SurvBal takes in a matrix consisting of the raw counts of the taxa for each subject in the study, a survival object from the R package “survival” [4] that contains survival/censoring times and indicators, as well as biomedical or demographic covariates that the users would like to adjust for in the analysis. The software is flexible as additional options are available throughout the model building and variable selection pipeline – from the pre-processing to the final selection of the microbial balance. Figure 1A provides an overview of the software, which we describe in greater detail below.

**Figure 1.**
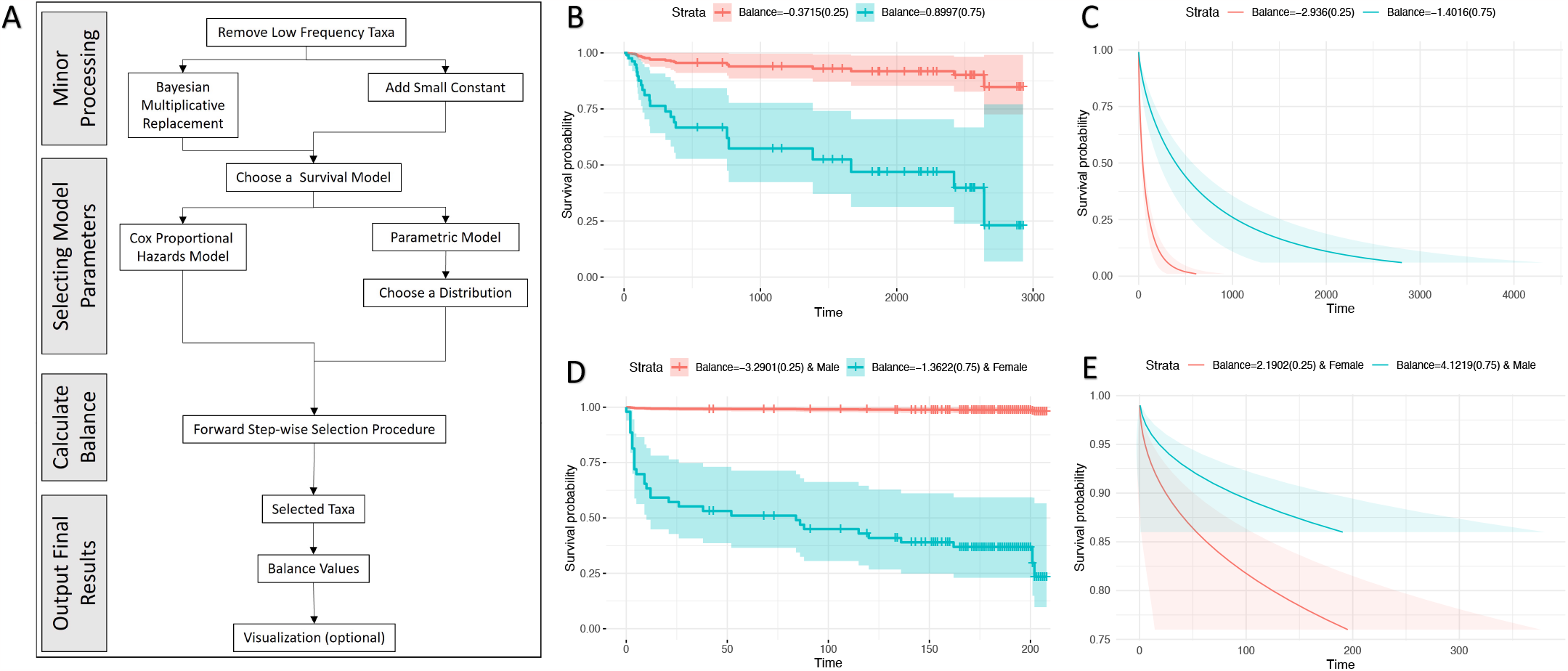
(**A**) Overview of SurvBal. Stratified time-to-event by a lower and a higher values of the selected balances of gut microbiome: HCT recipients, (**B**) OS under the Cox model, (**C**) Time-to-GvHD under the parametric survival model; kidney transplant recipients, (**D**) Time-to-*E*.*coli* bacteriuria under the Cox model, (**E**) Time-to-*Enterococcus* bacteriuria under the parametric survival model.

### 2.1 Review of Balance

Suppose we have ***X*** = (*X*_1_, *X*_2_, …, *X*_*k*_) as the microbial composition, where *X*_*i*_’s are relative abundances and summed up to 1. If we have a subset of bacteria that increase hazard or survival time, which is denoted by ***X***_+_, indexed by *I*_+_ and composed of *k*_+_ taxa, and another subset decreasing hazard or survival time, denoted by ***X***_*−*_, indexed by *I*_*−*_ and composed of *k*_*−*_ taxa, the global balance, or microbial signature, is defined as the normalized log ratio of the geometric means of the two groups,

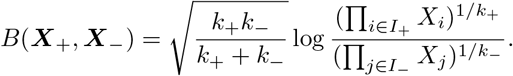

Equivalently, we can have

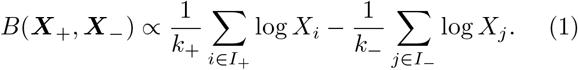

The log-contrast in (1) handles compositionality [5], accommodating the relative nature of microbiome abundances. Moreover, (1) is a special log-contrast, where the log differences are not included in a linear equation additively, but are combined together as a single variable [1].

### 2.2 Data Processing

Microbiome data are frequently sparse, i.e. contain excessive zeros, as some taxa are observed in only a subset of samples. Since SurvBal operates on the log scale of relative abundances, zeros need to be imputed. Here, after reading in the raw counts of the microbiome, SurvBal provides two commonly used options within the field – the geometric Bayesian multiplicative replacement (GBM) [6] (default), which is used in the original Selbal package, or adding a small constant (typically 0.5) to all values. The imputed counts will then be converted to relative abundance and taken log transformation. Furthermore, since some taxa are observed in only a small handful of samples, SurvBal allows users to provide a threshold such that taxa with prevalence below the threshold are dropped. This thresholding is particularly necessary when using GBM. Users can also process the microbial counts following their preferred routines before feeding the data into SurvBal, e.g., filtering out low-abundance taxa.

### 2.3 Choosing a Model

SurvBal offers two types of survival models, the Cox proportional hazards (default) and parametric survival models. The former is a semi-parametric model that assumes the hazards (instantaneous probability of failure) are proportional across stratification of covariates, while the latter assumes the survival time follows a particular distribution such as the Weibull distribution. Because a taxon increasing hazard naturally correlates with a shortened survival time, if a taxon appears in the nominator of the selected balance by the Cox model, likely, it will be added to the denominator of the balance by the parametric survival model. The scenario is just *likely* because the two models barely select the same set of taxa as they are fundamentally different in all aspects – from target variables, assumptions, to estimation algorithms. SurvBal allows both models to incorporate covariates such as crucial biomedical markers and demographic information. Furthermore, for parametric modeling, in addition to Weibull, all options in the original “survival” package can be specified, from exponential to log-logistic distributions.

### 2.4 Selecting the Global Balance

#### 2.4.1 Searching for the initial pair

First, SurvBal loops through all possible pairs of bacteria and determines the pair that is most correlated with the survival outcome. There are two combinations for each pair of taxa *r* and *s*, using the Cox model as an example, log *X*_*r*_ *−* log *X*_*s*_ assuming *r* is positively associated with risk and *s* is negatively associated with risk, and log *X*_*s*_*−*log*X*_*r*_ for the opposite scenario. Thus, essentially, SurvBal does an exhausted search checking the corresponding p-values of the 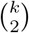 combinations, and picks the best pair with the minimal p-value while allocating the two bacteria into the nominator and denominate based on the sign of the pair’s associated coefficient. Suppose log *X*_*r*_*−*log *X*_*s*_ achieves the smallest p-value and its coefficient is positive, then the initial balance will be

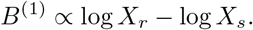

#### 2.4.2 Forward step-wise selection

Given *B*^(1)^, SurvBal searches through all the remaining taxa for the next one to include in the balance. For each taxon *t* (where *t* is not equal to *r* or *s*), we assess both ways to add it – adding it to the nominator or the denominator,

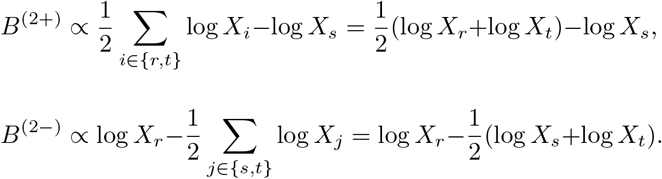

We calculate the p-values via the specified model regarding *B*^(2+)^ or *B*^(2*−*)^ as the balance variable. The procedure is repeated for each remaining taxon in turn. Taxon *t* that gives the smallest p-value among the (*k−*2) *×*2 choices will be selected and if the p-value corresponds to *B*^(2+)^ then taxon *t* goes into the numerator and if it corresponds to *B*^(2*−*)^ then taxon *t* goes into the denominator. We repeat the selection forward. Likewise, given *B*^(*u−*1)^, SurvBal determines *B*^(*u*)^ by picking one more taxon from the remaining pool excluding taxa already in *B*^(*u−*1)^ and adding it into the numerator or the denominator. The iteration terminates when it hits the stopping criteria – the p-value of the current balance is larger than a specified threshold (default=0.15). Optionally, if the users prefer sparser models, on top of the stopping p-value, they can turn on a sequential test that tests whether *B*^(*u*)^ is significantly different from *B*^(*u−*1)^ for the survival outcome in the specified model. If not, we stop adding more taxa. The significant level (default=0.05) of the sequential test can be tuned.

#### 2.4.3 Picking the optimal model

Although we stop adding taxa, we need to figure out which of the models in the selection sequence is the best one. In the previous steps, SurvBal keeps track of the models that include *B*^(1)^, *B*^(2)^, … as the balance variable. Then, in the final step, we can pick the one with the smallest p-value as the final model. This strategy is called the “minimum p-value” selection. Alternatively, we can pick the model of which the decrement of p-value to the next model is less than a specified threshold (default=15%) for the first time along the sequence. This is called the “minimum decrement of p-value” strategy, providing sparser models than “minimum p-value”, while the model fit could be similar. Therefore, for the sake of achieving moderate model complexity, “minimum decrement of p-value” is the default option.

### 2.5 Output and Visualization

In the output, SurvBal reports the selected global balance, including the selection sequence, the names of bacterial taxa included in the nominator and the denominator of the balance, and the balance value computed for each subject. It also presents the final survival model, the Cox proportional hazards or parametric survival model, with the selected balance and covariates specified by the users. Finally, it provides a visualization of the balance by examining predicted survival curves stratified by representative quantiles of the balance. SurvBal makes it flexible for users to explore the selected balance, allowing them to define interested quantile levels of the balance, and also the significance level for confidence intervals of those stratified survival curves.

## 3 Illustrating Examples

The first example is a study on graft-versus-host disease (GvHD) [7], a fatal complication of hematopoietic cell transplantation (HCT) and associated with the gut microbiome [8–11]. Here, we aim to find gut microbial signatures measured right after HCT to describe the overall survival (OS) and time to the GvHD. The processed data contained 63 recipients, and the 16S rRNA microbiome data was aggregated to the genus level with rare taxa (relative abundance *<* 0.01%) removed. The Cox proportional hazards model was used for analyzing OS and the parametric survival model assuming Weibull distribution was employed for time-to-GvHD. No covariates were adjusted and all other arguments were kept as the default options.

The global balance selected for OS by SurvBal includes 6 taxa increasing the risk of death 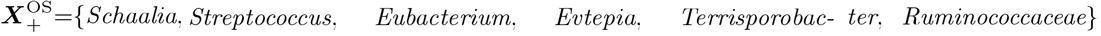, and 9 taxa decreasing the risk 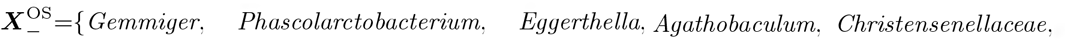 *Enterococcus, Pseudoflavonifractor/Clostridium, Collinsella, Romboutsia*}. Figure 1B presents the predicted OS curves. For recipients with a lower balance score (-0.37, the 1st quartile of the 63 recipients’ balances), which means that there are lower relative abundances of taxa in 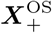 than in 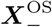, their life expectancy is significantly longer than those with a higher balance score (0.90, the 3rd quartile of the 63 recipients’s balances). The balance identified for time-to-GvHD comprises 4 taxa prolonging the time to the disease 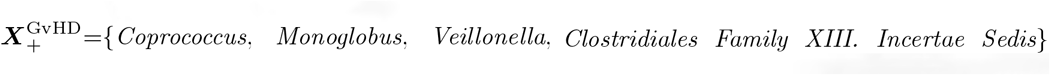, and 6 taxa shortening the time to the onset 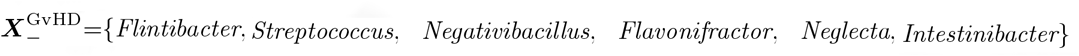, supported by the predicted time-to-GvHD curves in Figure 1C. The two balances for GvHD and mortality share *Streptococcus*, which shortens the time-to-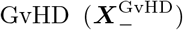 and increases the risk of death 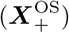. The positive association between *Streptococcus* and GvHD has been confirmed in previous HCT studies [12, 13]. With 63 patients only, some results obtained are contrary to literature such as *Enterococcus* was included in 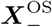 but has been linked with increased risk of GvHD [14]. Results will be further improved with more patients recruited.

The second example is a study on bacteriuria after kidney transplant [15, 16]. Bacteriuria is a common complication leading to significant morbidity, while modulating the gut microbiota could be a promising preventive intervention. Therefore, we would like to find the gut microbiome balance right after the surgery to predict the time to the onset of *Escherichia coli* and *Enterococcus* bacteriuria. The processed data contained 163 recipients, and the 16S rRNA microbiome data was aggregated to the genus level with common taxa (relative abundance *>* 1%) maintained. Cox and parametric models were used for *E*.*coli* and *Enterococcus* bacteriuria, respectively. Gender is a crucial risk factor for bacteriuria thus was adjusted in the analysis.

The balance determined for *E*.*coli* bacteriuria consists of 3 genera increasing the risk 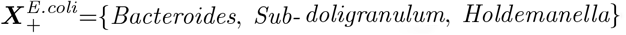, and 3 genera decreasing the risk 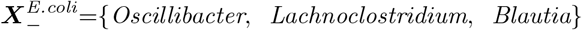 . The predicted time-to-*E*.*coli* bacteriuria is longer for recipients with lower relative abundances of taxa in 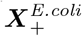 than in 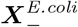, and the risk of *E*.*coli* bacteriuria is exaggerated within female recipients (Figure 1D). The balance selected for *Enterococcus* bacteriuria includes 1 genus extending the time to the disease 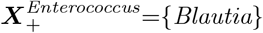, and 5 genera expediting the onset 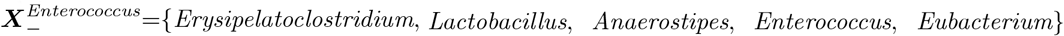. SurvBal successfully identified *Enterococcus* as a factor for promoting *Enterococcus* bacteriuria, which is consistent with prior work [16]. The predicted time-to *Enterococcus* bacteriuria is systematically shortened with a lower balance score (Figure 1E). *Blautia* is associated with both preventing the onset of *Enterococcus* 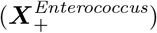 and decreasing the risk of *E*.*coli* bacteriuria 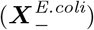. It is a common commensal bacteria in the gut and is associated with the production of butyrate, a short-chain fatty acid (SCFA) [17]. Prior work [18] has shown that SCFAs can inhibit the growth of *E. coli* via intra-cellular acidification, suggesting a competitive role of this bacteria against *E. coli* in the gut.

## 4 Conclusion

We have developed the R package SurvBal to enable the analysis of compositional balances in microbiome profiling studies with censored time-to-event outcomes. The software allows the users to flexibly incorporate covariates and choose their preferred model type and other options. We expect that the selected global balance will further guide clinical investigations into the microbial markers, which can be modulated to improve the time-to-event outcomes.

## Data Availability

GvHD data (de-identified for patient confidentiality) is available upon request. KTx data (sequencing and deidentified clinical data) is available in dbGaP (accession number: phs001879.v1.p1) and the access needs local institutional review board approval.

## Acknowledgments

We thank Dr. Manikkam Suthanthiran for providing the support for the KTx dataset.

